# Applying optimal control theory to complex epidemiological models to inform real-world disease management

**DOI:** 10.1101/405746

**Authors:** E.H. Bussell, C.E. Dangerfield, C.A. Gilligan, N.J. Cunniffe

**Author notes:** **Corresponding Author:** Dr. Nik Cunniffe. **Twitter:** @nikcunniffe.

## Abstract

Mathematical models provide a rational basis to inform how, where and when to control disease. Assuming an accurate spatially-explicit simulation model can be fitted to spread data, it is straightforward to use it to test the performance of a range of management strategies. However, the typical complexity of simulation models and the vast set of possible controls mean that only a small subset of all possible strategies can ever be tested. An alternative approach – optimal control theory – allows the very best control to be identified unambiguously. However, the complexity of the underpinning mathematics means that disease models used to identify this optimum must be very simple. We highlight two frameworks for bridging the gap between detailed epidemic simulations and optimal control theory: open-loop and model predictive control. Both these frameworks approximate a simulation model with a simpler model more amenable to mathematical analysis. Using an illustrative example model we show the benefits of using feedback control, in which the approximation and control are updated as the epidemic progresses. Our work illustrates a new methodology to allow the insights of optimal control theory to inform practical disease management strategies, with the potential for application to diseases of plants, animals and humans.

## 1 Introduction

Mathematical modelling plays an increasingly important role in informing policy and management decisions concerning invading diseases [1, 2]. However, model-based identification of effective and cost-efficient controls can be difficult, particularly when models include highly detailed representations of disease transmission processes. There is a variety of mathematical tools for designing optimal strategies, but no standard for putting the results from mathematically motivated simplifications into practice. An open question is how to incorporate enough realism into a model to allow accurate predictions of the impact of control measures, whilst ensuring that the truly optimal strategy can still be identified [3]. In this paper we identify the difficulties – as well as potential solutions – in achieving a practically useful optimal strategy, highlighting the potential roles of open loop and model predictive control by way of a simple example.

### Realistic landscape-scale modelling

The optimisation of epidemic interventions involves determining the management strategy that minimises impacts of disease. This minimisation can be difficult when resources are limited and there are economic costs associated with both control measures and disease. Methods that simulate the expected course of an epidemic and explicitly model effects of interventions allow us to quantify the potential impact of a given strategy [4]. These simulation models accurately capture the dynamics of the real system and so have become important tools in designing intervention strategies to inform policy decisions. Examples include vaccination policies for human papillomavirus in the UK [5, 6], livestock culling policies for foot-and-mouth disease [7, 8] including post-hoc vaccination optimisation [9, 10], and optimal host removal radii for tree diseases of citrus [11–13] and sudden oak death [14].

Various complexities of disease dynamics, for example spatial heterogeneities and inherent individual differences in susceptibility and pathogen transmission (risk structure), have been shown to be important determinants of patterns and rates of epidemic spread [15–17]. To ensure accurate epidemic predictions, these factors must be included in simulation models designed to aid decision making. However, inclusion of these factors typically results in highly complex models with many possible control measures, making optimisation computationally infeasible when interventions can be combined, and particularly when control measures can also vary over time, in space or according to disease risk [18]. For most simulation models the only viable option is then to use the model to evaluate a small subset of plausible strategies that remain fixed during the epidemic, potentially scanning over a single parameter such as a culling radius. We shall refer to this approach as ‘Strategy Testing’. Using this approach makes it difficult to have high confidence in the best-performing strategy, since with no framework for choosing it, the set of strategies under test is likely to be biased. Further to this, as the set to test cannot span the entire space of control options, it is unlikely that the true optimum will be found.

### Optimal control of epidemiological models

Many mathematical techniques exist for characterising the true optimal control for a disease, such as equilibrium or final size analysis, depending on the system being analysed [15]. We here focus on optimising time-varying control of dynamical systems, for which optimal control theory (OCT) is widely used [19]. The controls considered in this paper are mathematical representations of possible disease interventions. By analysing a set of equations describing the disease dynamics, OCT can mathematically characterise the optimal control and provide insight into the underlying dynamics, without the repeated simulation required to optimise simulation models. However, because of the underlying mathematical complexity, little progress can be made with OCT unless the underpinning models for disease spread are highly simplified. Early work in OCT focussed on optimal levels of vaccination and treatment [20], with extensions to consider further interventions including quarantine, screening, and health-promotion campaigns appearing later [21]. Disease models can also be coupled with economic effects [22–24], and within OCT this has been used to balance multiple costs, such as surveillance and control [25], or prophylactic versus reactive treatment [26].

The optimal strategies identified by OCT can be very complex, often specifying controls that switch strategies at specific times during the course of an epidemic. The added complexity of these switching controls can significantly improve disease management when tested on a spatially explicit model, but can perform poorly if the exact time of the switch is not known [27], for example when parameters are not known precisely. This demonstrates that numerical results from OCT are not always directly applicable to the real world because of additional complexities in real systems. It is also unclear however, how insight from OCT alone could be translated into practical advice. To move towards robust strategies that could be used practically, more recent work has focussed on including additional features and heterogeneities into the models used in OCT, in particular spatial dynamics. Space is usually only included to a limited extent, for example by using metapopulation models (e.g. [28, 29]), or partial differential equations (e.g. [30]) to optimise spatial strategies, so whether the heterogeneities added are sufficient to identify robust and practical control strategies remains an open question.

### Moving towards practical control

Despite finding the mathematically optimal control strategy, major simplifications to the system as modelled are required to allow progress to be made using OCT. It is therefore often unclear how these strategies would perform if adopted by policy makers. On the other hand, models with sufficient realism to inform policy directly are often impossible to fully optimise. Therefore, a framework is needed to couple the results of OCT with the realism required in policy making and in simulation type models. The question is then how should we make practical use of OCT?

In §2 we describe two methods from control systems engineering for applying OCT results, and compare these versus Strategy Testing using a simple illustrative model in §3. We seek to answer how, under current computational constraints, results from OCT can be applied whilst maintaining the realism required for practical application.

## 2 Applying optimal control to realistic systems

Outside of epidemiology, OCT has had wider use on approximate models of complex systems. A recent study reviews the use of OCT for agent-based models (ABMs) [31], a type of model that simulates the individual behaviour of autonomous agents. An *et al*. [31] suggest the use of a model that approximates the dynamics of the ABM, designed to be simple enough to allow mathematical analysis of the optimal control. A suitable approximate model is chosen and fitted either to real data, or to synthetic data from the ABM. The OCT results from the approximating model are then mapped onto the ABM to be tested: a process referred to as ‘lifting’, which could equally well apply to the detailed epidemic simulation models considered in this paper. We now describe two possible frameworks from control systems engineering for making use of this control lifting approach.

### Open-loop control

The first method is the simplest application of control lifting, and the framework implicitly suggested by An *et al*. [31]. Control is optimised on the approximate model once using the initial conditions of the simulation model, and this control is lifted to the simulator and used for the full simulation run time (figure 1). Crucially, any time variation in this control is fixed at the very start of the simulation, and so does not incorporate any feedback. It is therefore referred to as ‘open-loop’ control, as it is fully specified by the simulation initial conditions and the trajectory predicted by the approximate model. Use in epidemiology is uncommon, although Clarke *et al*. [32] use OCT in an approximate model to find optimal levels of Chlamydia screening and contact tracing which are then mapped onto a network simulation.

**Figure 1:**
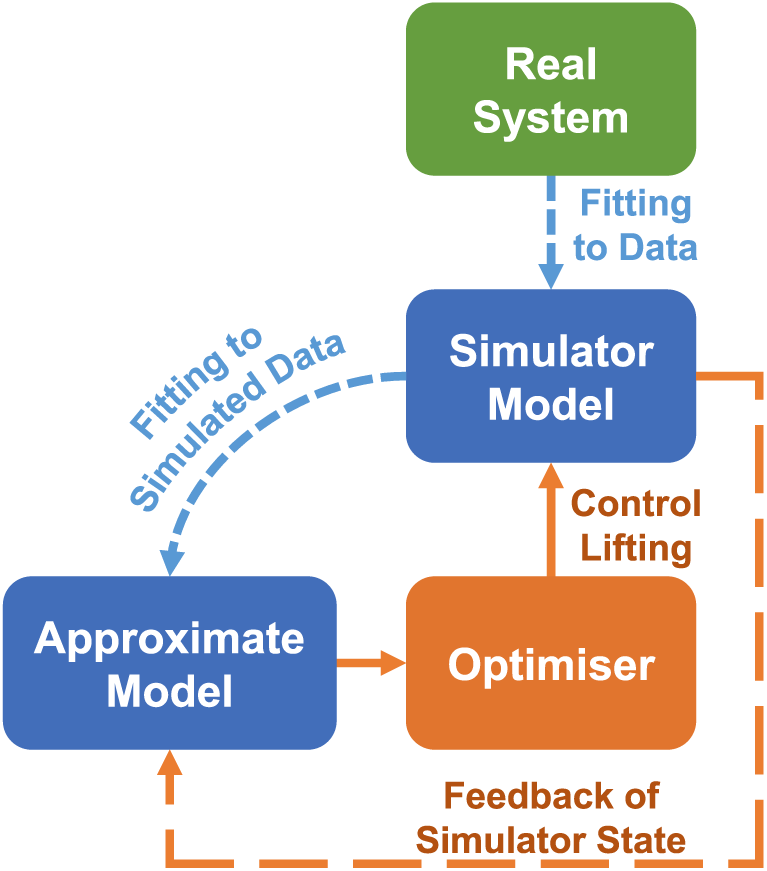
Open-loop and model predictive control (MPC). The model hierarchy is shown, with optimised controls from the approximate model directly lifted to the simulation model. The real system is in green, the models and fitting processes are in blue, and the control framework is in orange. Without the orange dashed feedback loop, this is open-loop control. MPC resets the state of the approximate model at regular update steps, before re-optimising and lifting controls to the simulation model until the next update time.

### Model predictive control

Open-loop control requires the approximate model to remain accurate over a long time scale. Since simulation models typically include many heterogeneities that must be omitted from the approximate model, systematic deviations between the simulation and approximate predicted trajectories for disease progress are highly likely. Model predictive control (MPC) is an optimisation technique incorporating system feedback [33, 34]. At regular update times the values of the state variables in the approximate model are reset to match the conditions in the simulation at that time. The control is then re-optimised and the new control strategy is used going forward in the simulation to the next update time. This corresponds to a series of open-loop problems solved at regular update steps (figure 1). Model predictive control can therefore take into account perturbations in the disease progress trajectory caused by heterogeneities omitted from the approximate model.

Model predictive control has had some use within the epidemiological literature, the majority being for control of drug applications for single individuals rather than control of epidemics at the population level. Examples include finding management strategies for HIV that are robust to measurement noise and modelling errors [35, 36], and control of insulin delivery in patients with diabetes [37]. These studies highlight the benefits of MPC for robust control, i.e. control that remains effective despite system perturbations. However, only one study concentrates on epidemic management [38], and that does not explicitly test the feedback control on simulations.

## 3 Optimising strategies on an illustrative network model

### Methods

To demonstrate open-loop and MPC for epidemic management we use a stochastic SIR network model including host demography and risk structure. The model is deliberately kept simple to show how the underpinning idea is broadly applicable across human, animal and plant diseases. Whilst the model and its parameters are arbitrary we use it to represent a scenario in which a simulation model has already been fitted to a real disease system; the network model is therefore used here as a proxy for a potentially very detailed simulation model.

### Simulation Model

In our model, infection spreads stochastically across a network of nodes that are clustered into three distinct regions (figure 2a). Each node contains a host population stratified into high and low risk groups. The infection can spread between individuals within nodes and between connected nodes. The net rate of infection of risk group *r* in node *i* is given by:

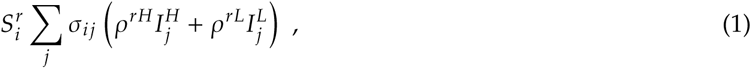

where *S* and *I* are numbers of susceptible and infected hosts respectively, subscripts identify the node, and superscripts specify high (*H*) or low (*L*) risk group. The sum is over all connected nodes including the focal node itself, with pairwise connectivities given by *σ_ij_*, and risk structure given by *ρ*. Full details of the model are given in the supplementary material. Although not limited to these applications, the model in Equation 1 could represent crop or livestock diseases spreading through farms, or sexually transmitted infections spreading through towns, cities or countries.

**Figure 2:**
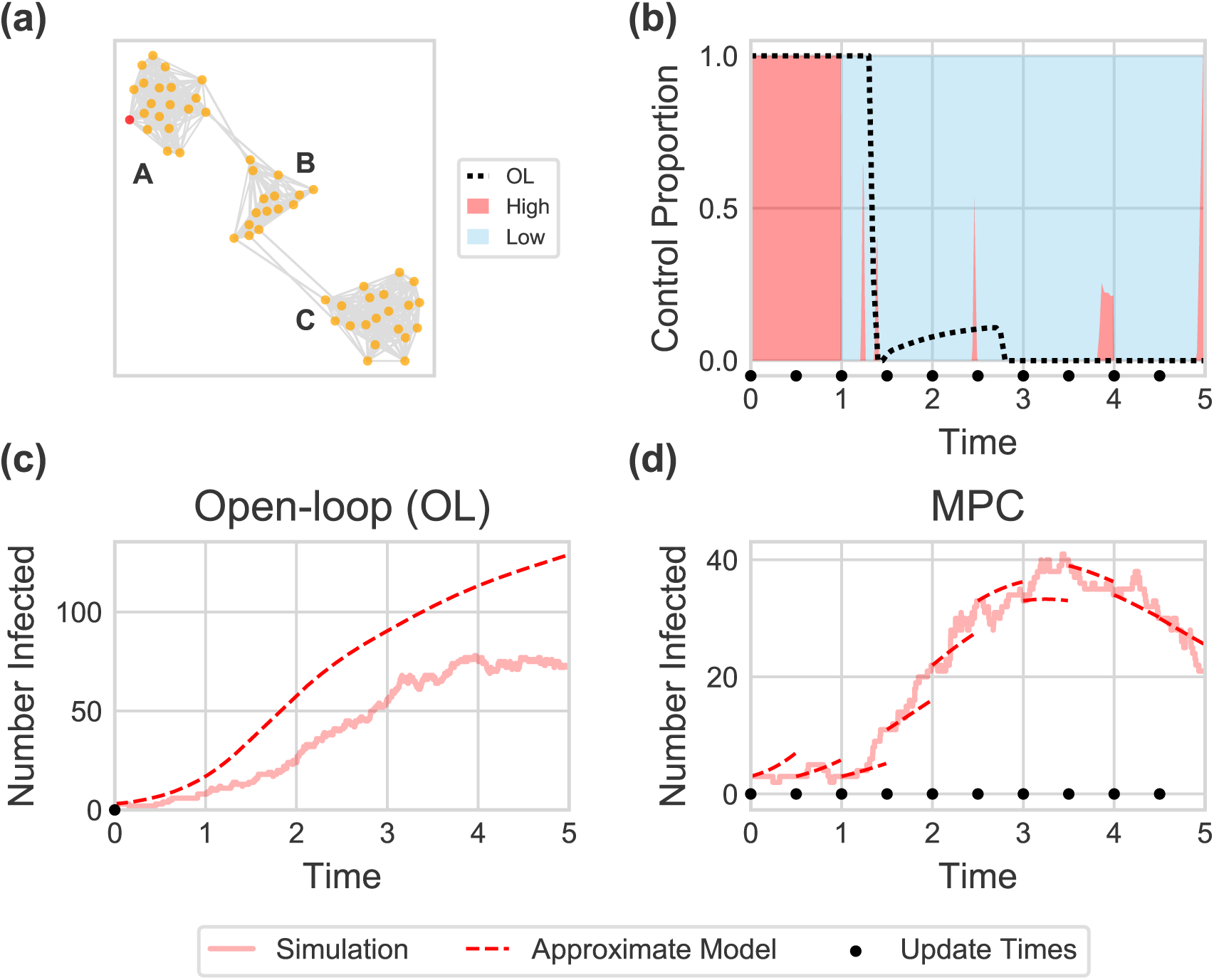
**(a)** shows the network used for the illustrative simulation model, including region labels. The epidemic is seeded in the red node in region A, and can spread between connected nodes (grey lines). In **(b)** the control allocation is shown for a single space based MPC run, with the corresponding open-loop allocation indicated by the black dotted line. **(c)** shows the total number of infected individuals under a single run of space based open-loop control. Control is based on the prediction of the approximate model starting from the initial conditions. **(d)** shows the number of infected individuals in the simulation and space based approximate model corresponding to the MPC control carried out in **(b)**. Here the prediction is reset to match the simulation at every update step (0.5 time units) and the control is re-optimised. By taking account of differences in the number of infected individuals compared with those predicted at the initial time, MPC gives better predictions of the simulation state as well as improved control when compared to open-loop control (note different y axis scales).

Mass vaccination is the only intervention we consider, with the potential to target based on both risk group and region but randomised across host infection status (i.e. susceptibles cannot be preferentially vaccinated). Logistical and economic constraints are included through a maximum total vaccination rate (*η*_max_) that can be divided between risk groups and regions. Within each group susceptibles are vaccinated at rate: *f η*_max_*S*/*N*, where *f* is the proportion of control allocated to that group, and *N* is the total group population.

Optimal allocation of the vaccination resources minimises an objective function *J* representing the disease burden of the epidemic across all infected hosts over the simulation time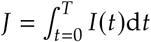. In common with the particular control we consider and the risk and spatial structures, this simple choice of objective function was made merely to illustrate our methods, but the framework generalises immediately to more complex settings.

### Approximate Models

Exhaustive optimisation of control using the simulation model, across space, risk group and time, is clearly infeasible. We consider two different deterministic approximate models of the simulator to assess the best level of approximation. The first model is purely risk structured, factoring out all spatial information and leaving one high risk and one low risk population group. The second approximate model is more complex, in as much as it is also risk structured, but additionally includes a first approximation to the host spatial structure by dividing the host population into three large regions. Spatial dynamics are included between but not within the three regions, maintaining enough simplicity to obtain optimal control results. This could represent, for example, optimising control at the country level, but not at the regional level. We will refer to this model as the spatial approximate model. A single set of parameters is fitted for each model to data from an ensemble of simulation model runs. We then test which of the two approximate models is the more useful for control optimisation. Full details of the approximate models, fitting and optimisation procedures are given in the supplementary material.

### Control Scenarios

We test six different control scenarios, which compare Strategy Testing (scenarios 1 and 2) with open-loop and MPC applied to both of our approximate models (scenarios 3 to 6):
1. ‘High’: exclusively vaccinate high risk individuals
2. ‘Split’: partition control resources between high and low risk groups based on an optimisation performed in advance
3. ‘Risk OL’: open-loop control on the risk based approximate model
4. ‘Risk MPC’: MPC on the risk based approximate model
5. ‘Space OL’: open-loop control on the spatial approximate model
6. ‘Space MPC’: MPC on the spatial approximate model

The optimal constant allocation for the ‘Split’ strategy was found by selecting the proportionate allocation to each risk group which minimised the mean disease cost in a set of simulation realisations. (supplementary material text and figure S8). Figures 2c and 2d illustrate the open-loop and MPC strategies, showing the model resetting at regular intervals in MPC.

## Results

The risk based optimisation results in a strategy that initially vaccinates high risk individuals, before switching priorities and treating the more populous low risk group almost exclusively (figure 2b). This same switch is seen in the optimal controls from the spatial approximate model, but a number of spatial switches are also seen (figure S9). The spatial strategies are therefore much more complex than the risk based controls. Comparing disease costs shows that incorporating greater realism, through a more complex approximate model as well as by using MPC, allows for improved management of the epidemic (figures 3 and S10). Of the constant ‘user-defined’ strategies, splitting control between risk groups is more effective than just vaccinating the high risk group. The optimal allocation to the high risk group is 63% of vaccination resources, with the rest used to vaccinate low risk individuals. Using the risk-based approximate model gives an improvement over either of these strategies, although there is little difference between open-loop and MPC (see below). Adding space into the approximate model improves control further, leading to the smallest epidemic costs when the MPC framework is used.

**Figure 3:**
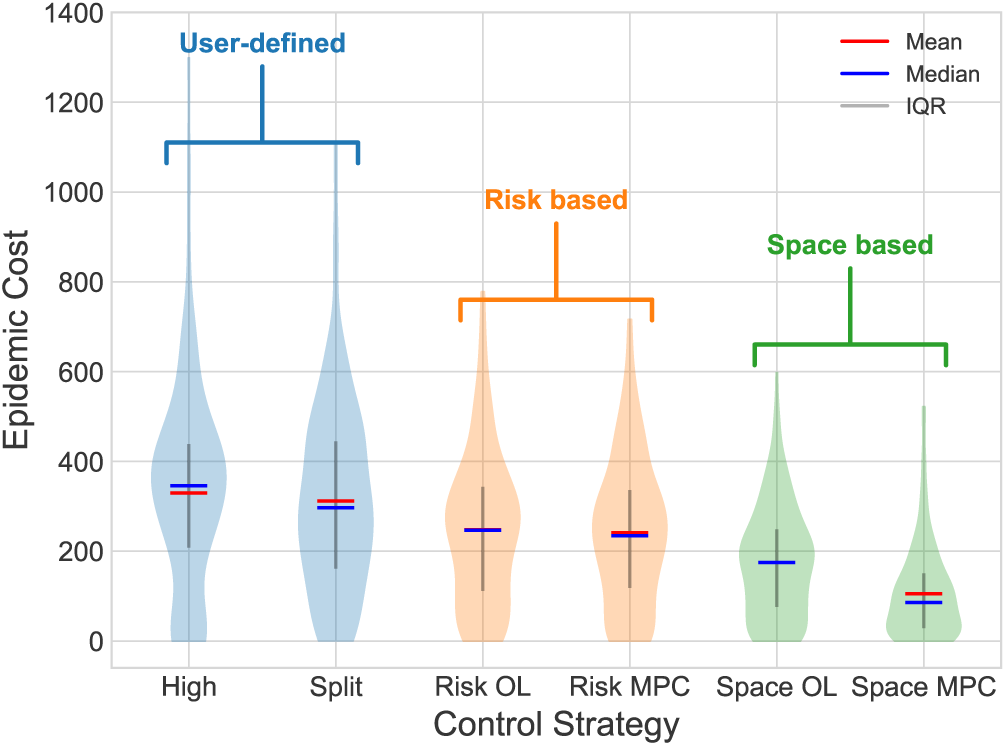
Results of different control optimisation schemes on the example simulation model. Spatial MPC performs best, showing an improvement over both open-loop and user-defined strategies.

The illustrative model demonstrates the management improvements that can be achieved by combining OCT with both open-loop and MPC. The key results of the OCT analyses are the control switching times. Using the switching controls from either approximate model with open-loop control gives lower epidemic costs than the naively chosen ‘user-defined’ strategies. Feedback allows the MPC controllers to re-evaluate the timing of these switches during the epidemic, and potentially include additional switches, to match the exact trajectory of the current realisation (figures 2b–d).

In the risk based strategies there is little difference between open-loop and MPC. This is because the precise timing of the switch from high to low risk group vaccination does not significantly affect the epidemic cost (supplementary figure S11). The timings of disease introduction into regions B and C are highly variable between simulation runs (supplementary figure S2). The potential for additional switches in the spatial approximate model gives more flexibility for the MPC controller to respond to this variability, and so spatial MPC shows a significant improvement over open-loop. Extending the objective to include a cost associated with switching strategy would allow a more detailed assessment of the practicality of implementing these complex strategies, although to provide a simple demonstration of our key ideas we have not pursued this here.

In the spatial case, open-loop performs worse than MPC as it is unable to adapt to perturbations in the dynamics. The control must then rely on the model predicted spread, necessarily an approximation to the actual trajectory. The performance of the control is closely linked to the accuracy of this approximation, raising the important issue of choosing the approximate model. In our example, spatial dynamics are clearly important because of the timing of spread between regions, and so the more informed controls of the spatial model outperform the risk based strategies.

## 4 Discussion and outstanding questions

Our results show that the choice of approximate model affects the performance of both open-loop and MPC strategies. Here we have found a suitable approximate model in an ad hoc manner, but a key challenge for the future is to develop a more formal method for choosing the most appropriate approximate model. A more accurate model will give better predictions, and hence control that is closer to the true optimum, but this must be balanced with added complexity and optimisation constraints. One difficulty in doing this is that it is not always clear where the boundary of mathematical or computational feasibility is, and so how complex the model can be made in practice. It is also difficult to mathematically determine, in a systematic way, which aspects of the dynamics are important to capture accurately. This key issue must be considered though, since the implications relate directly to applications in the real world.

Practical disease control requires surveys of the real system to assess the state of the epidemic. Both open-loop and MPC optimise control using predictions of the future dynamics, making them both feed-forward controllers. The approximate model underlying these frameworks allows more informed decisions between surveys, resulting in control that is closer to the true optimum. Accurate predictions can avoid continuous or very frequent surveys which may be expensive or logistically challenging. The repeated updates of MPC improve these predictions but will each require surveillance of the real system, so the frequency of updates must be chosen so as to balance improved knowledge of the system with any surveillance constraints.

In addition to this feed-forward control, MPC incorporates a feedback loop not included in open-loop strategies. The feedback mechanism, included through the update steps, allows for control that copes better with uncertainty, i.e. it is more robust. As shown by [35], MPC can still successfully control a system when measurements of that system are inaccurate, or the parameters in the simple approximate model are incorrect. In our case this allowed it to perform better on the complex simulation model, where additional heterogeneities were present. Open-loop strategies are not robust to these perturbations so may not perform well in practice. For certain systems and parameters the open-loop strategy may in fact perform worse than naive constant control [28]. We also see this for certain parameter sets in the illustrative model. It would be easy to dismiss the OCT results as not useful for policy, but our results show that by using feedback, these results can still be beneficial.

In this paper we have focussed on a top-down approach, finding robust, practically-applicable strategies by making use of OCT to optimise simulation models. Equally, many studies analyse dynamical systems using OCT without simulation models. These studies rarely consider practical application of the resulting optimal controls, and so may benefit from a framework for testing the results. With this bottom-up approach, a system for testing the results on realistic systems is vital to ensure that these results are robust to additional realism. Using an MPC framework as considered here could be one way in which OCT researchers could demonstrate the potential impact of their work to a wider audience.

We have assumed throughout that an accurate simulation model of the real system in question can be built, and that a single set of parameters can be fitted for the chosen deterministic approximate model. In reality there may be considerable uncertainty in parameters for the simulator so fitting a single deterministic model may be challenging. A question for future study would be how to handle these uncertainties, perhaps also incorporating improved knowledge of parameters as the simulation proceeds [39]. A final challenge for optimal control more generally relates to the choice of objective function. The strategies found by OCT are highly dependent on the exact form of the objective function, and so more research is needed into how to quantify the balancing of very different costs, for example treatment costs and disease burden [28]. In human disease, cost-effectiveness analyses are usually based on quality adjusted life years (QALYs) [40], and a maximum economic cost per QALY gained. A similar concept could perhaps be used for plant and animal diseases, including calculations of yield losses [41] as well as effects on welfare, biodiversity and tourism for example [42].

## 5 Conclusions

Simulation models that capture the complex dynamics and uncertainties necessary for informing policy have significant limitations when designing optimal time-dependent management strategies. Optimal control theory is a powerful mathematical approach for characterising these interventions, but results have proved difficult to translate into effective policy advice. Increased focus on how optimal control can be used and translated to policy makers would allow for more robust decisions on disease management, incorporating greater epidemiological understanding.

Lifting of controls from OCT to realistic simulations as in [31] goes some way to addressing this problem, but the differences in model structure between the simulation and approximate models is likely to limit the practical performance of the control measures. In this paper we show that incorporating feedback can alleviate this problem and help lead to robust and practical control strategies. Whilst these techniques may be able to transfer optimal control results to more realistic simulations and so to practical application, it does raise the issue of communicability of results. With complex feedback strategies between two models, one complex in structure and the other mathematically complex, the overall result is no longer simple to explain. How applicable these strategies can ever be, will then be determined by the reliability of, and trust in, the simulation model, and its ability to establish the benefit of a more complex intervention strategy.

Overall, there is clearly a benefit to policy and management decisions from making use of OCT results. We suggest that potential policies, including OCT results, can and should be tested on realistic simulations, and that MPC is an effective platform for practical applications of OCT.

## Additional Information

## Acknowledgements

We thank Andrew Craig, Eleftherios Avramidis and Hola Adrakey for helpful discussions.

## Data Accessibility

All code and generated data are available at https://github.com/ehbussell/Bussell2018Model.

## Authors’ Contributions

E.H.B., C.E.D. and N.J.C. designed the study, E.H.B. conducted the analysis and wrote the initial draft of the manuscript. All authors contributed to data interpretation, manuscript editing and discussion.

